# Frontocentral brain networks amplify action-oriented temporal prediction errors

**DOI:** 10.1101/2025.08.07.669061

**Authors:** Davide Sometti, Ileana Rossetti, Cornelius Schwarz, Markus Siegel, Christoph Braun, Antonino Greco

**Affiliations:** Graduate Training Center of Neuroscience, International Max Plank Research School, University of Tübingen, Tübingen, Germany; Department of Neural Dynamics and Magnetoencephalography, Hertie Institute for Clinical Brain Research, University of Tübingen, Tübingen, Germany; Werner Reichardt Centre for Integrative Neuroscience, University of Tübingen, Tübingen, Germany; MEG Center, University of Tübingen, Tübingen, Germany; Center for Bionic Intelligence Tübingen Stuttgart (BITS), Tübingen, Germany; Department of Psychology, University of Milano-Bicocca, Milan, Italy; Milan Centre for Neuroscience (NeuroMi), University of Milano-Bicocca, Milan, Italy; Department N3: Neurorehabilitation, Neuroprosthetics, Neurotechnology, Hertie Institute for Clinical Brain Research, University of Tübingen, Tübingen, Germany; German Center for Mental Health (DZPG), Tübingen, Germany

## Abstract

Successful sensorimotor behavior depends on anticipating when a tactile event will occur and translating it into rapid action, yet it remains unclear how action requirements shape temporal prediction in the human cortex. Here, we combined Magnetoencephalography (MEG), multivariate decoding and computational modeling to contrast temporally jittered finger stimulations under passive versus action-oriented conditions. We characterized a distributed frontal sensorimotor network that was linked to fast tactile-motor associations and predicted reaction-time variability. We found that trial-by-trial deviations from expected stimulus timing modulated behavior, with stronger frontocentral prediction-error encoding when an overt response was required. Temporal prediction errors were primarily encoded in a motor rather than sensory reference frame. These results reveal that action demands dynamically amplify temporal error signaling in sensorimotor circuits, in line with predictive coding frameworks postulating that the brain preferentially encodes task-relevant and goal-directed sensory prediction errors.

## Introduction

Somatosensory-motor interaction is essential for flexible and efficient behavior^1,2^, and becomes especially critical when rapid responses to external events are required. Consider, for example, a tennis player sensing the subtle vibrations through the racket as the ball makes contact, those fleeting tactile cues must be translated almost instantaneously into precisely timed muscle activations to return a fast serve or powerful groundstroke. Even for highly practiced actions, successful performance relies on the brain’s ability to predict when that tactile input will arrive and to prepare the appropriate motor output in advance.

Contemporary predictive coding theories of brain function suggest that the brain is constantly generating and updating predictions about incoming sensory input, minimizing the mismatch between expectations and actual events^3–12^. Within this framework, temporal expectations are a particular form of prediction, tuning neural activity so that the system is optimally ready at anticipating events^13–17^.

Prior work indicates that the brain can extract sensory regularities even when they carry no immediate behavioural relevance^7,10,18^ and that prediction error signals are amplified for events tied to task demands^19,20^. However, it remains unclear whether this amplification occurs in sensorimotor tasks when an overt action is required, and which neural mechanisms support the encoding of these prediction errors.

To address these questions, we combined Magnetoencephalography (MEG) with multivariate decoding and computational modeling. We characterized the spatiotemporal dynamics of temporal prediction error encoding in the human brain during a tactile-motor association task. Participants received temporally jittered brief pulses on one finger and either simply perceived them (stimulus-only blocks) or responded as fast as possible with a finger movement (stimulus-response blocks). This paradigm enabled us to investigate how the brain anticipates and responds to behaviorally relevant, temporally variable tactile input and how temporal expectations about that input are encoded and linked to upcoming motor output, when one must act versus merely passively perceive. Furthermore, we investigated whether prediction errors were encoded in sensory or motor reference frames.

## Results

We analyzed data from 11 human participants performing a tactile-motor experiment, with a total of 4 different tasks performed during the MEG measurement (Fig. 1a). In 2 of the tasks, namely stimulus-only (S-O) and response-only (R-O), stimulus and response were independent to each other and serve as a baseline: participants had to either passively receive a tactile stimulation on the index fingertip or to press a button with the thumb. The other 2 tasks, referred to as stimulus-response (S-R) tasks, combined the two previous conditions by linking the tactile stimulation to the motor response. Specifically, participants had to react as fast as possible to the tactile stimulation by responding with a button press. Stimulated and responding hands could be the same (ipsilateral: S_Lx_R_Lx_ and S_Rx_R_Rx_) or different (contralateral: S_Lx_R_Rx_ and S_Rx_R_Lx_). All tasks were performed with both hands for a total of 8 experimental conditions, each condition consisting of 130 trials. Importantly, the interstimulus interval (ISI) between each stimulation in S-O and S-R conditions uniformly varied between 2.5 and 3 seconds, allowing participants to form temporal expectations. Behavioral compliance to S-R tasks was controlled through reaction time (RT, Fig. 1b). No differences were found across conditions (all *p* > 0.054; all Cohen’s *d* < 0.386), median reaction times (with IQR) were as follows: S_Lx_R_Lx_ = 0.203 s (0.192 - 0.230 s); S_Lx_R_Rx_ = 0.191 s (0.165 - 0.214 s); S_Rx_R_Rx_ = 0.209 s (0.186 - 0.266 s); S_Rx_R_Lx_ = 0.213 s (0.179 - 0.250).

**Figure 1.**
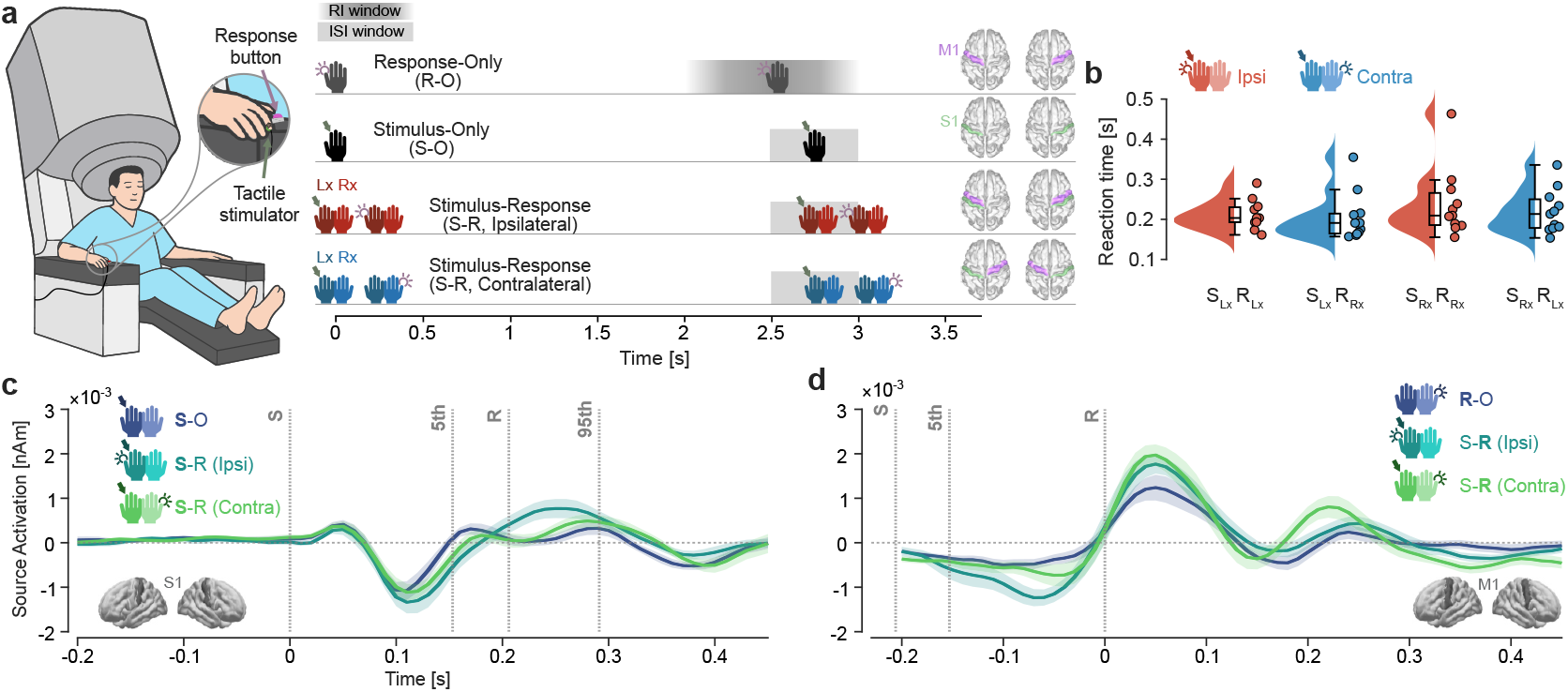
Experimental design, behavioral performance and sensorimotor responses. **a**. Illustration of the experimental setting in the MEG scanner. Participants performed four tasks: response-only (R-O), stimulus-only (S-O), and two stimulus-response (S-R) tasks with either ipsilateral (Ipsi) or contralateral (Contra) stimulus-response association. Each condition was repeated for both hands. Shaded areas represent response intervals (RI) and inter-stimulus intervals (ISI) temporal windows. On the right, schematic illustration of the primary sensory (S1) and motor (M1) areas involved in each condition. **b**. Reaction time (RT) distributions across the four S-R conditions. Each dot represents individual participants. **c, d**. Time courses of source activation in primary somatosensory (S1) and motor (M1) cortices across S-O, S-R (Ipsi), and S-R (Contra) conditions, averaged across hemispheres. Activity profiles are time-lock aligned to stimulus and response events. Shaded areas indicate the standard error of the mean (SEM). Vertical dotted lines represent the stimulus onset (S), the response onset (R) and the 5^th^ and 95^th^ percentile of the temporal onset distribution.

We started analyzing the neural data by reconstructing brain activity at the source level using linearly constrained minimum variance beamforming^21^ and parcellated activity into 72 brain regions according to the Desikan-Killiany scheme^22^. We plotted the average time course of source activity across primary somatosensory (S1) and motor (M1) cortices, aligning trials to stimulus onset or the response (Fig. 1c-d). In S1, we observed a clear peak around 50 ms after stimulus onset, followed by one deflection around 100 ms, while in M1, a readiness potential was observable before the movement and was followed by two peaks around 50 and 200 ms after response onset.

### Neural dynamics during tactile-motor association

We investigated the neural patterns underlying tactile-motor associations using multivariate decoding^23^. Classification analysis was performed over time using a surface-based searchlight approach to compare S-O vs. S-R conditions aligned to stimulus onset (Fig. 2a-c) and R-O vs. S-R aligned to response onset (Fig. 2b-d; see Supplementary Fig. 1 for analyses within individual conditions). We found that both Ipsi and Contra S-R conditions were well decodable against S-O (Fig.2a; Ipsi: *p* < 0.001 cluster-corrected, mean *d* = 2.645; Contra: *p* < 0.001 cluster-corrected, mean *d* = 2.799) and RO (Fig.2b; Ipsi: *p* < 0.001 cluster-corrected, mean *d* = 2.782; Contra: *p* < 0.001 cluster-corrected, mean *d* = 2.540). Decodability started around 100 ms prior to stimulation, reached peak accuracy around response time, and further decreased gradually. No significant difference in decoding accuracy was found between Ipsi and Contra conditions (p > 0.05 cluster-corrected).

**Figure 2.**
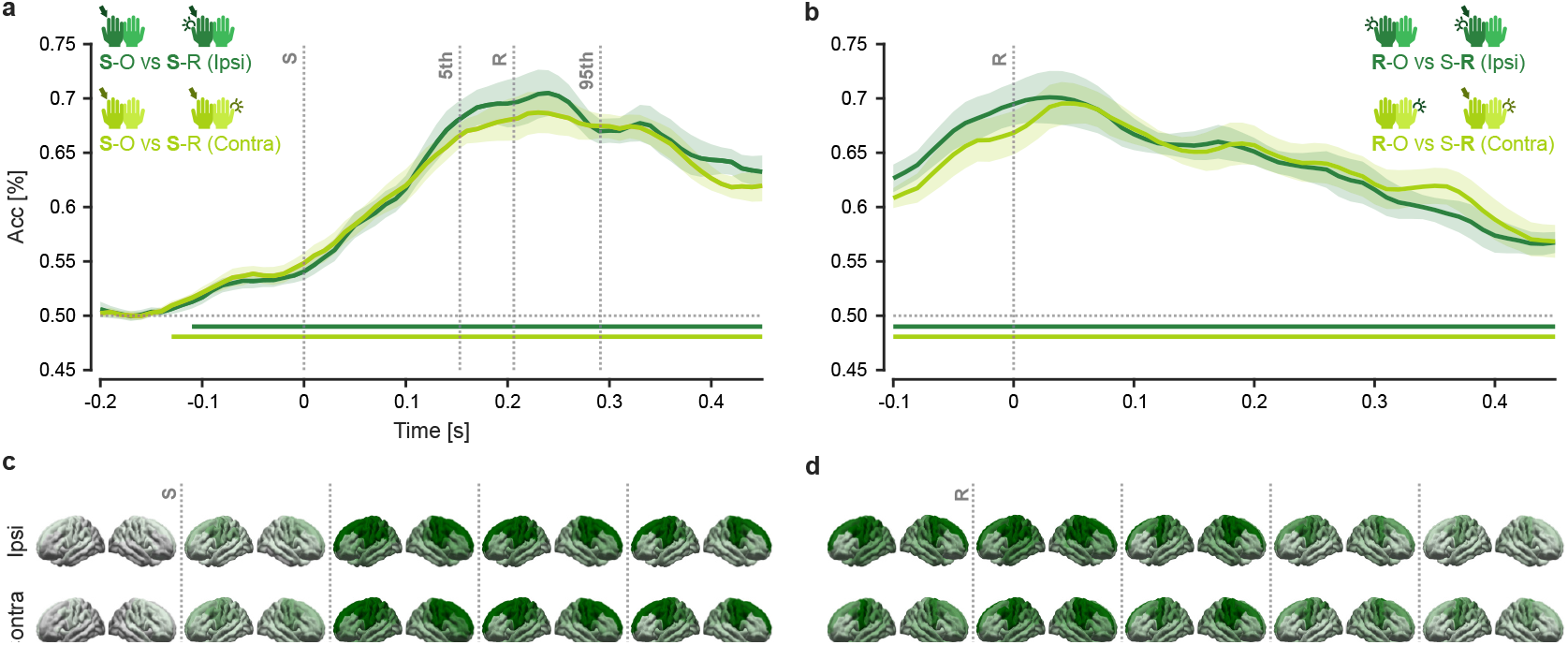
Fronto-central cortical patterns underlie S-R association. Time-resolved multivariate decoding accuracy (Acc) discriminating S-R (ipsilateral and contralateral) conditions from stimulus-only (S-O, panel **a**.) and response-only (R-O, panel **b**.) baseline conditions. Shaded areas indicate SEM, and horizontal lines indicate statistical significance (p < 0.05) above chance level. Vertical dotted lines represent the stimulus onset (S), the response onset (R) and the 5^th^ and 95^th^ percentile of the temporal response onset distribution. Below, cortical surface maps from searchlight decoding analyses illustrate the spatial dynamics of S-R versus S-O (**c**.) and R-O (**d**.) conditions. Statistical significance was determined by cluster based one-tail permutation test (p < 0.05) and highlighted with opacity across the cortical surface.

This classification analysis, repeated in a searchlight fashion across the cortical surface, highlighted the contribution of a broad cluster of fronto-central areas during the association of stimulus and response. This, in line with the time-resolved decoding, showed how the contribution of the cluster to the decodability increased over time, starting from before stimulus onset (Fig.2c; across all time-windows, Ipsi: all *p* < 0.001 cluster-corrected, 1.004 < mean *d* < 2.383; Contra: all *p* < 0.001 cluster-corrected, 1.005 < mean *d* < 2.429) and increasing until the motor response before gradually decreasing after completion of the task (Fig.2d; across all timewindows, Ipsi: all *p* < 0.001 cluster-corrected, 1.680 < mean *d* < 2.33; Contra: all *p* < 0.001 cluster-corrected, 1.663 < mean *d* < 1.976).

To determine whether the progressive emergence of a fronto-central network of brain areas during stimulus–response (S–R) association reflected functional relevance to task performance, we used a multivariate linear regression approach to predict RT from neural data aligned to stimulus onset in the S–R conditions. Time-resolved decoding revealed that RTs were significantly predicted in both Ipsi and Contra conditions from peristimulus onset (Fig. 3a; Ipsi: p < 0.001 cluster-corrected, mean d = 1.524; Contra: first cluster −80-0 ms, p = 0.038 cluster-corrected, mean d = 0.797, second cluster 50-210 ms, p < 0.001 cluster-corrected, mean d = 2.224).

**Figure 3.**
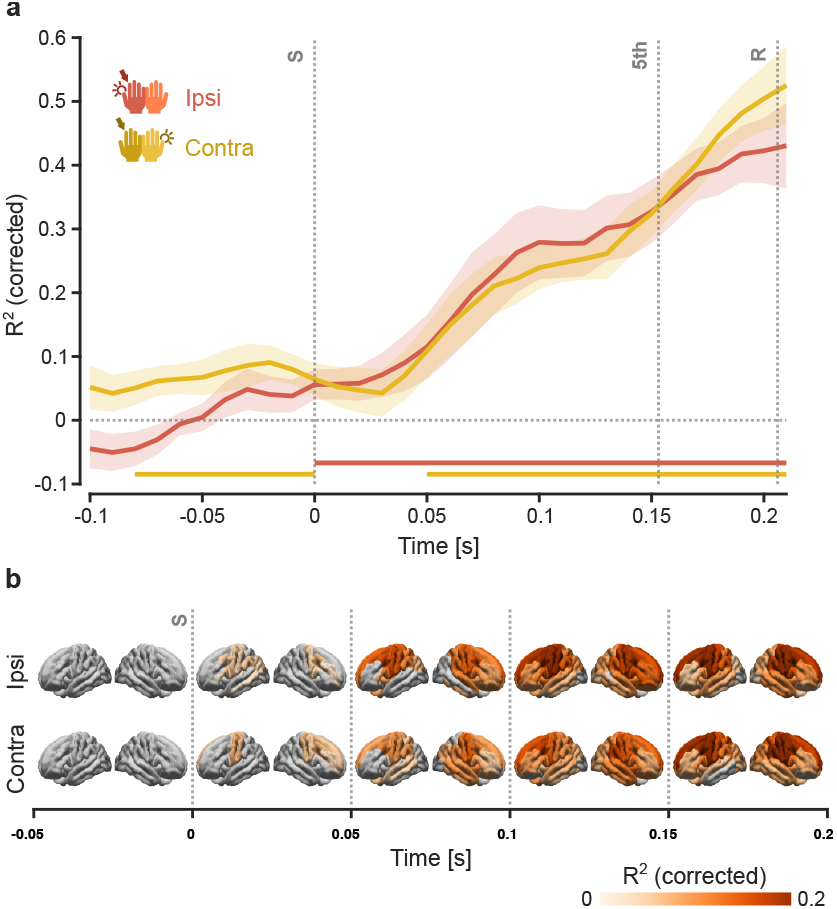
Fronto-central cortical patterns predict reaction time during stimulus-response association. **a**. Time-resolved multivariate regression (R^²^ corrected) of brain activity predicting reaction time (RT) for ipsilateral and contralateral stimulus-response (S-R) conditions, aligned to stimulus onset. Shaded areas indicate SEM, and horizontal lines indicate statistical significance above chance prediction performance (p < 0.05), determined with a one-tailed cluster-based permutation test. **b**. Searchlight regression across the cortical surface highlights the spatial distribution of predictive activity over time. Significance was determined by cluster-based one-tail permutation test (p < 0.05) and highlighted with opacity across the cortical surface. Vertical dotted lines represent the stimulus onset (S), the response onset (R) and the 5^th^ percentile of the temporal response onset distribution.

When assessing the spatial distribution of this effect using a searchlight approach, we observed in both conditions a fronto-central network emerging around stimulus onset (Fig. 3b), consistent with classification findings. Significant spatial clusters were observed in the temporal window from 0 to 200 ms, in both Ipsi (all p < 0.007 cluster-corrected, 0.778 < mean d < 1.327) and Contra (all p < 0.046 cluster-corrected, 0.748 < mean d < 1.496) conditions.

### Amplified action-oriented error encoding in fronto-central activity

We next focused on our key question, whether temporal prediction errors are differentially encoded in the peristimulus neural response depending on whether participants must act upon or merely perceive the tactile stimulus. To test this, we modeled each trial’s temporal expectation using a rolling average, where the expected onset on each trial was computed as the average ISI over a window of *ω* previous trials. We defined a prediction error as the difference between the actual and expected ISI, reasoning that participants would respond more slowly when the stimulus arrived earlier than expected (negative error) and more quickly when it arrived later than expected (positive error). By correlating the trial-by-trial error trajectory for each participant with RTs, we observed a robust negative relationship (Fig. 4a), in line with our expectation. We performed model comparison to identify the best *ω* parameter and selected the window size (*ω* = 2) that yielded the strongest negative correlation between error and RT across participants. Using this window size parameter, we extracted the prediction error trajectories (Fig. 4b) from the model and split RTs following positive versus negative expectation errors and confirmed that RTs were significantly faster following positive errors in both conditions (Fig. 4c, Ipsi: *p* < 0.001, *d* = 1.453; Contra: *p* = 0.002, *d* = 1.447).

**Figure 4.**
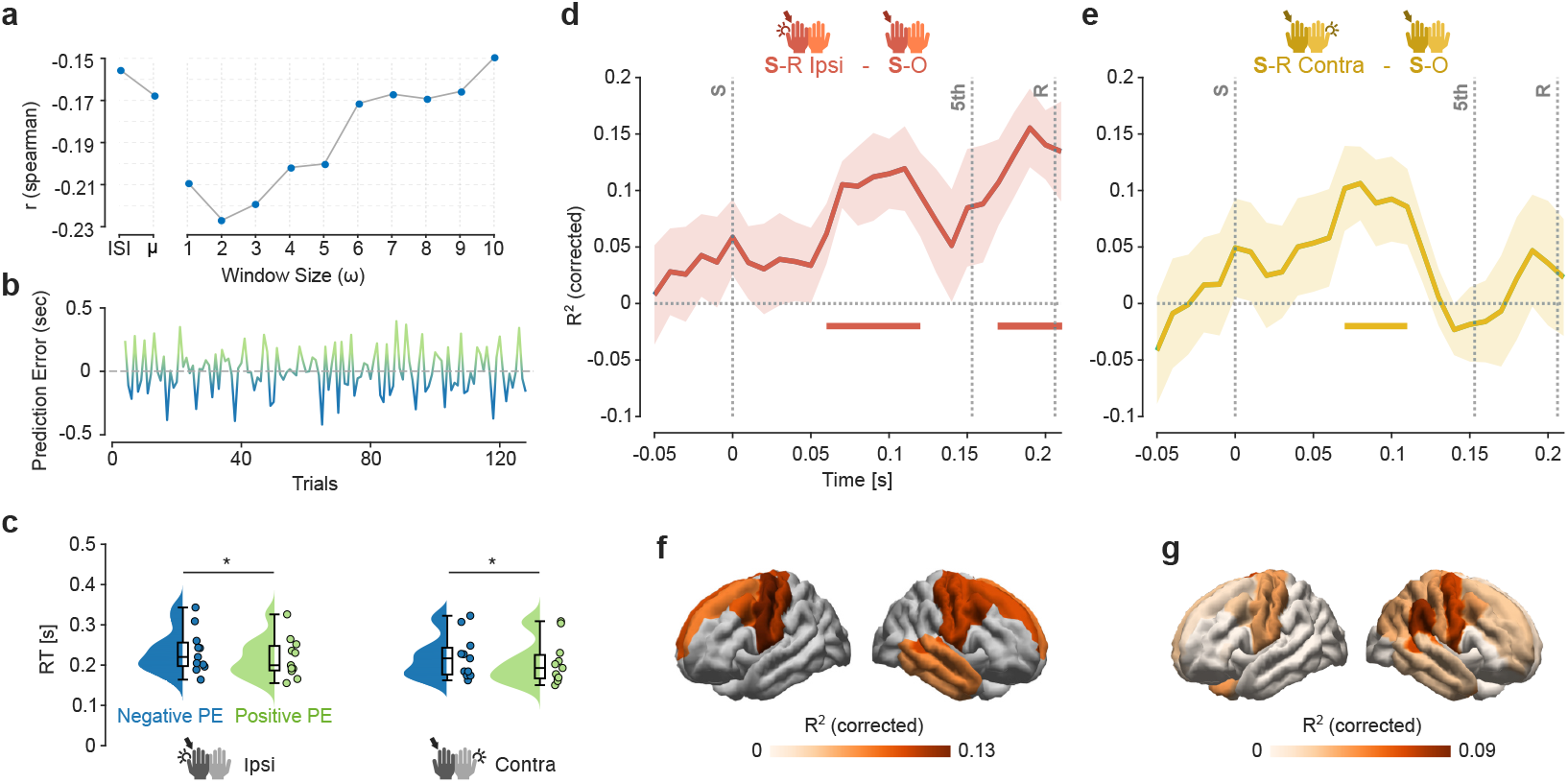
Neural encoding of temporal prediction errors during tactile–motor association. **a**. Spearman correlation between prediction error and RTs across window sizes (*ω*) on pooled data across S-R conditions. **b**. Lineplot showing an exemplar prediction error trajectory across trials using the winning model (*ω*=2), color-coded by the errors being positive or negative. **c**. Raincloud plots depicting RTs split by positive versus negative prediction errors. Each dot represents individual participants while horizontal bars indicate statistical significance (*p* < 0.05). Lineplots showing the timeresolved multivariate regression (R^²^, corrected) model performance of peri-stimulus neural activity predicting trial-wise prediction error in S-R Ipsi (**d**.) and S-R Contra (**e**.) conditions minus the S-O condition. Shaded areas represent SEM. Horizontal bars indicate statistical significance (cluster-based permutation test, p < 0.05). Vertical dotted lines represent the stimulus onset (S), the response onset (R) and the 5^th^ percentile of the temporal response onset distribution. Below, surface-based searchlight regression showing the cortical distribution of the effects found in the time-resolved analysis, in S-R Ipsi (**f**.) and S-R Contra (**g**.) conditions minus the S-O condition.

We next asked when and where neural activity would selectively reflect prediction errors under action demands, by contrasting time-resolved multivariate regressions of stimulus-aligned MEG data against the trial-by-trial error signal in the stimulus–response (S–R) versus stimulus–only (S–O) conditions. This direct comparison isolates activity related to response-dependent temporal expectation encoding, since both models use the prediction-error trajectory but only the S–R blocks require a motor response. We first tested the encoding of prediction error against chance and found significant effects only in S-R conditions but not in S-O (Supplementary Fig. 2a; S-O, p > 0.05, cluster corrected, 2b; S-R Ipsi, *p* = 0.0024 cluster-corrected, mean *d* = 0.928, 2c; S-R Contra, *p* = 0.023 cluster-corrected, mean *d* = 0.819).

Remarkably, by directly contrasting S-R and S-O conditions, we found that when an action was required the prediction error was amplified compared to when participants only received passive stimulation, with the encoding of prediction error showing its peak around 100 ms post-stimulus. Specifically, we identified two significant temporal clusters in the Ipsi condition (Fig. 4d; 60–120 ms, *p* = 0.0093 cluster-corrected, mean *d* = 0.981; 170–210 ms, *p* = 0.017 cluster-corrected, mean *d* = 1.111), and one in the Contra condition (Fig. 4e; 70–110 ms, *p* = 0.0469 cluster-corrected, mean *d* = 0.803). Searchlight analysis, focusing on the earliest temporal window per condition, revealed in the Ipsi a significant cluster across fronto-central and right temporal cortical areas (Fig. 4f; *p* = 0.002 cluster-corrected, mean *d* = 0.745). In the Contra condition, a similar fronto-central and right temporo-parietal distribution appeared but this effect did not reach statistical significance (Fig. 4g; *p* > 0.05 cluster corrected).

### Temporal prediction errors are encoded in motor rather than stimulus space

We conducted a control analysis to ensure our effects were not driven by the motor response itself. We first averaged all S-R trials time-locked to the button-press to create a motor-evoked template. We then subtracted this template from each individual S-R trial (aligned at its own response time), yielding a set of response-subtracted S-R trials (Fig. 5a).

**Figure 5.**
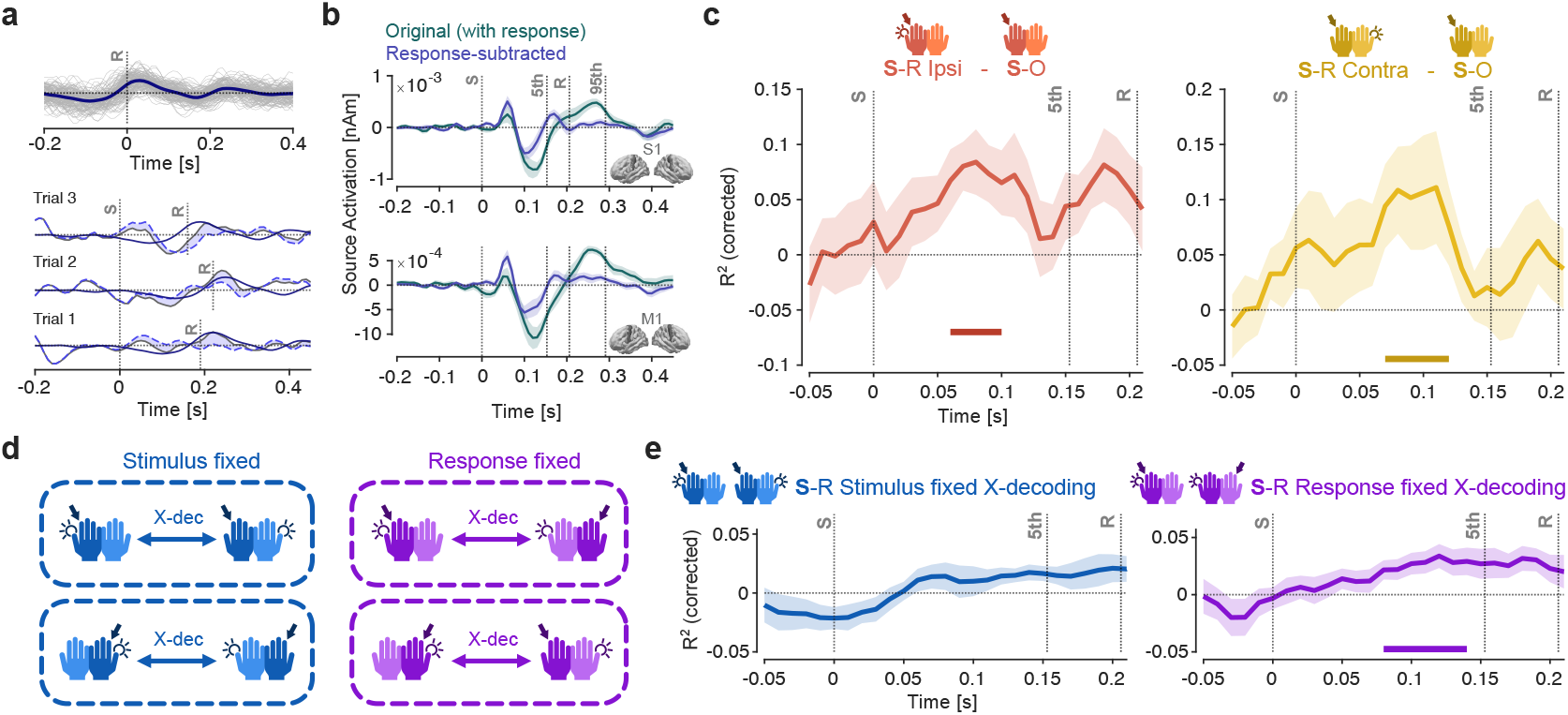
Temporal prediction errors are encoded in motor reference frame. **a**. Schematic illustration of the procedure for obtaining response-subtracted data. Top, example of motor template evoked activity by averaging S-R trials aligned to the response. Below, exemplar trials showing the motor template signal (dark blue line aligned to the response onset), original signal (grey solid line), the response-subtracted signal (blue dotted line) and its difference as shaded blue area. **b**. Lineplots showing the group-average activity in S1 and M1 with original (with response) and response-subtracted data. Shaded areas represent SEM, while vertical dotted lines represent the stimulus onset (S), the response onset (R) and the 5^th^ and 95^th^ percentile of the temporal response onset distribution. **c**. Lineplots showing the time-resolved multivariate regression (R^²^, corrected) model performance of peri-stimulus neural activity predicting trial-wise prediction error in S-R Ipsi and S-R Contra conditions minus the S-O condition using response-subtracted data. Shaded areas represent SEM. Horizontal bars indicate statistical significance (cluster-based permutation test, p < 0.05). **d**. Schematic illustration of the cross-decoding (X-dec) procedure implemented to test pattern generalization of neural mechanisms encoding temporal prediction errors, testing stimulus and response fixed conditions. **e**. Lineplots indicating the cross-decoding in S-R stimulus and response fixed conditions using response-subtracted data. Shaded areas represent SEM. Horizontal bars indicate statistical significance (cluster-based permutation test, p < 0.05), while vertical dotted lines represent the stimulus onset (S), the response onset (R) and the 5^th^ percentile of the temporal response onset distribution

When these adjusted trials were plotted in the S1 and M1 ROIs, the source-reconstructed activity around the response window was markedly altered, confirming that our subtraction procedure successfully isolated and removed the motor component (Fig. 5b). We repeated the same time-resolved multivariate regression analysis, which confirmed our previous results. Specifically, we found that S-R conditions showed error encoding by contrasting against chance (Supplementary Fig. 3a; Ipsi, 30–210 ms, *p* < 0.0001 cluster-corrected, mean *d* = 1.392; 3b; Contra, −40–20 ms, *p* = 0.006 cluster-corrected, mean *d* = 1.149, 80-140 ms, *p* = 0.004 cluster-corrected, mean *d* = 1.233) and, crucially, against S-O condition (Fig. 5c; Ipsi 60–100 ms, *p* = 0.030 cluster-corrected, mean *d* = 1.159; Contra, 70–120 ms, *p* = 0.028 cluster-corrected, mean *d* = 1.111). In sum, these findings suggest that temporal prediction errors are stronger encoded in fronto-central cortices when an overt response is required compared to passive stimulation.

Finally, we investigated whether error signals were represented in a sensory versus motor reference frame. To address this, we performed the same time-resolved multivariate regression analysis as above, but in a cross-decoding fashion^24^ (Fig. 5d). We trained the regression model on one S-R condition and testing it on another such that either the motor response was held constant, but the sensory input differed across conditions (e.g., train on S_Lx_R_Rx_ and test on S_Rx_R_Rx_) or vice versa. Remarkably, we found significant pattern generalization only when the motor output was identical (Fig. 5e; 80–140 ms, *p* = 0.006 cluster-corrected, mean *d* = 1.125), but not when the sensory input was held constant (*p* > 0.05 cluster-corrected), suggesting that temporal prediction errors were primarily encoded in a motor rather than sensory reference frame.

## Discussion

Our study provides insights into the neural encoding mechanisms of temporal prediction errors related to action demands during tactile-motor associations. We characterized a frontal-sensorimotor network that exhibited anticipatory, predictive and reactive control dynamics linked to behavioral performance.

Neural spatiotemporal patterns remained consistent when decoding was performed separately for each S–R pairing, indicating that the observed effects are not linked to condition pooling but rather reflect a generalized interhemispheric mechanism for tactile-to-motor transformation. Accordingly, previous fMRI studies have revealed limb-independent frontoparietal representations of hand actions during movement planning ^25^ and shown that movement type can be decoded from a widespread bilateral network of motor and somatosensory areas during execution^26^. In our study, however, the motor response required no action selection, suggesting these neural patterns instead reflect broad anticipatory sensorimotor readiness^27,28^, driven by predictive signals^29–31^ and supported by frontal top-down preparation and inhibitory control^32–34^. We found that a similar frontocentral network of brain areas predicts reaction time variability. Previous results indicate that prefrontal regions exert an attention based top-down modulation over somatosensory cortex during tactile discrimination^35–37^. In the present task, similar mechanisms may have facilitated the rapid sensorimotor transformation required to convert a brief, temporally unpredictable tactile event into a speeded motor response^38^.

Our observation that prediction-error signals are amplified in fronto-central circuits when an overt response is required resonates with a growing body of work showing that the brain’s predictive machinery is not purely automatic but is flexibly tuned by task relevance^19,39,20,40–42^. Auksztulewicz and colleagues demonstrated in a visual MEG paradigm that both prediction and prediction-error signals, indexed respectively by beta/alpha and gamma-band activity, are markedly stronger when spatial or temporal cues are behaviorally relevant versus irrelevant^20^. Similarly, Stokes et al. used EEG to show that statistically identical regularities are coded more robustly for target-relevant predictions than for non-target contingencies, with enhanced evoked responses to predicted targets emerging around 200 ms post-stimulus only when those predictions directly supported task goals^19^. In line with these findings, our direct contrast between stimulus–response and stimulus–only blocks reveal that imposing an action requirement enhances temporal-prediction signals in somatosensory-motor networks enabling rapid behavioral adjustments. Under a predictive-coding framework, this dynamic weighting according to task demands can be understood as a mechanism for allocating processing resources to the most behaviorally pertinent predictions and errors, ensuring that sensory processing and motor planning remain tightly coordinated in service of adaptive action ^43–46^.

Our finding that temporal prediction error encoding generalizes only across conditions sharing the same motor output indicates that the human cortex rapidly re-codes timing deviations into a motor reference frame, underscoring the central role of action in human cognition. Contemporary theories of embodied and enactive cognition posit that cognition is inseparable from action, with perception arising through our engagement with environmental affordances^47^, shaped by the rules governing sensorimotor interactions^48^ and constituting a form of enactive sense-making grounded in bodily dynamics^49^. Our findings also accord well with the active inference account within the predictive-coding framework, which further underscores action’s primacy in cognition^43– 45,50^. Rather than passively updating beliefs, the brain actively samples the world by issuing motor predictions that minimize surprise. This motor-centric encoding likely confers several functional advantages. By aligning prediction-error signals with the effector systems that will implement corrective behavior, the brain can enact swift adaptations without the need for costly coordinate transforms. Moreover, maintaining errors in a common motor frame may support the flexibility to generalize across different sensory contexts^51^.

In conclusion, our findings show that action demands amplify temporal prediction errors in frontocentral networks and are represented in a motor reference frame. Our results support predictive coding theories that posit action as the primary scaffold for encoding task-relevant sensory prediction errors.

## Acknowledgements

This study was supported by the European Research Council (ERC; https://erc.europa.eu/, CoG 864491 to M.S.) and by the German Research Foundation (DFG; https://www.dfg.de/, projects 276693517 (SFB 1233; to M.S.) and SI 1332/6-1 (SPP 2041; to M.S.)).

## Author contributions

D.S.: conceptualization, software, methodology, investigation, formal analysis, validation, data curation, visualization, writing - original draft, writing - review and editing.

I.R.: conceptualization, investigation, writing - review and editing

C.S.: conceptualization, writing review and editing.

M.S.: conceptualization, methodology, supervision, resources, project administration, funding acquisition, writing - review and editing.

C.B.: conceptualization, methodology, supervision, project administration, writing - review and editing.

A.G.: conceptualization, methodology, supervision, project administration, software, formal analysis, visualization, writing - original draft, writing - review and editing.

## Materials and Methods

### Participants

Twelve healthy participants (7 male) between 21 and 39 years old (mean ± SD age: 27.83 ± 5.46 years) participated in the study. One participant was excluded due to very low behavioral performance, resulting in a final sample of 11 participants (see the preprocessing subsection for detail). All participants were right-handed, provided informed consent prior to the experimental session, and received monetary compensation after completing the study. The protocol was in accordance with the Helsinki Declaration (2012) and approved by the ethics board of the University of Tübingen Medical School (protocol number 539/2020BO).

### Task

The experiment consisted of 3 different experimental tasks: stimulation-only (S-O), response-only (R-O) and stimulus-response (S-R) tasks. During the S-O task subjects passively received tactile stimulation on the index fingertip delivered through a pneumatic stimulator. The non-magnetic device featured an inflatable membrane clipped to the finger to ensure stable contact with the skin. It was briefly inflated by pressurized air delivered from an external compressor through a small plastic tube. Each inflation of the membrane produced a clear tactile sensation, exerting a small pressure (~1.6 N) on the fingertip. A total of 130 stimulations of 50 ms duration each was delivered in one single condition. Importantly, the interstimulus interval (ISI) between each stimulation uniformly varied between 2.5 and 3 seconds.

In the R-O condition participants were instructed to repeatedly press an optical button every 2 to 3 seconds in a self-paced manner using the thumb for a total of 130 presses. Finally, the S-R condition consisted in a reaction time (RT) task where the subject was asked to respond as fast as possible to the tactile stimulation of the index fingertip by pressing a button with the thumb. The ISI between the stimulation was the same as for the S-O condition while the response depended on the reaction time of the participant.

Both left and right hand were involved in the experiment leading to a total of 8 experimental conditions: 2 for the S-O, 2 for the R-O and 4 for the S-R since all combinations of the stimulation and response sides were considered. Out of the stimulus-response conditions, 2 were ipsilateral with stimulation and response on the same side (e.g. left index fingertip stimulation, left thumb response) and 2 were contralateral with stimulus and response at different hands (e.g. left index fingertip stimulation, right thumb response). The sequence of the conditions was randomized across subjects.

Throughout the entire experiment, participants remained comfortably seated in a relaxed posture inside the magnetically shielded measurement chamber, with their arms resting on the chair’s armrests. They were asked to avoid any muscle contraction not associated with the task. White noise presented via a pair of compatible earplugs was used to mask any acoustic cue emitted by the pneumatic stimulator.

### MEG data acquisition and preprocessing

MEG data were recorded using a whole-head 275 sensors CTF MEG system (VSM Medtech, Port Coquitlam, Canada) installed in a magnetically shielded room (Vacuumschmelze, Hanau, Germany). MEG signals were sampled at 585.94 Hz.

All the analyses were carried out using MATLAB (version 2021b). Preprocessing, source reconstruction and statistics were computed using FieldTrip (version 20211118) ^52^. Multivariate classification and regression analysis were performed using the MVPA-Light toolbox ^53^.

We started analyzing behavioral performance during S-R conditions. First, we analyzed the performance within the single subject, to ensure a stable performance we rejected trials whose RT was lower or higher than 3 scalar median absolute deviation (MAD) ^54^. Then, using the same metric, we analyzed the performance across subjects, excluding one participant whose performance was consistently slower in all 4 S-R conditions.

Continuous data were filtered between 1 and 30 Hz by applying a sixth-order Butterworth high-pass and low-pass filter respectively. Prior to segmentation we corrected for a 40 ms delay between the recorded trigger signal and the pneumatic stimulation onset. S-O and R-O conditions were segmented in trials of 650 ms duration, from −200 to 450 ms relative to the trigger onset. S-R data were segmented from −200 ms before the stimulus onset to 450 ms after the motor response, each trial length varied depending on the reaction time.

Cleaning of the neuromagnetic data was performed using a semi-automatic procedure: trials and channels were visually inspected for artifacts and removed whenever exceeding a cutoff threshold of variance < 10^−25^ T^2^. Independent component analysis ^55^ was applied to remove non-cortical physiological activity (eye movement, heartbeat, muscle contraction) using the logistic infomax ICA algorithm ^56^ to decompose the signal. Topography and temporal profile of the identified components were visually inspected, and any components of artifactual origin were discarded.

### Source reconstruction

Linearly constrained minimum variance (LCMV) beamform^21^ was used to reconstruct source activity. This spatial filtering method imposes a unit-gain constraint, by passing the signal at a specific source location without attenuation, while minimizing the contribution from sources at any other location, under the assumption that the underlying sources are temporally uncorrelated ^57^. For each subject, we computed the individual forward model (leadfield) between sources and sensors. This was done using a single shell head model (Nolte 2003). We co-registered the brain template *fsaverage* (Fischl et al. 1999) with the MEG sensors based on the individual’s head position. The head position was registered during the measurement using three head localization coils attached to the nasion and left/right preauricular points.

Leadfield and data covariance matrix were used to compute the spatial filter for the source reconstruction. Two spatial filters were computed to reconstruct single-trial sensor-level activity. The first filter, aligned to stimulus onset, combined S-O and S-R conditions using a covariance matrix spanning 0 to 180 ms post-stimulation. The second filter, aligned to response onset, combined R-O and S-R conditions with a covariance matrix covering −200 to 400 ms around response time. Trial counts were equalized across conditions to ensure balanced filter weights. Both covariance matrices were regularized (λ = 5%) for numerical stability^57^. Trials were then resampled to 100 Hz, projected to source space, applying the two filters separately to generate stimulus- and response-aligned datasets. Finally, source space was parcellated into 72 brain areas using the Desikan-Killiany atlas^22^.

### Classification analysis

Linear discriminant analysis (LDA) was selected as a classifier using accuracy (ACC) as a performance metric. To ensure robust and balanced model evaluation, we implemented a 10-fold stratified cross-validation procedure with two repetitions, integrating random oversampling within each training fold to address class imbalances while maintaining representative class distributions across folds. To reduce overfitting, the optimal shrinkage parameter for the regularization of the training set covariance matrix was calculated, using the Ledoit-Wolf method^58^.

Classification was performed on source reconstructed data. S-R conditions were split in ipsilateral, when stimulus and response involved the same hand, and contralateral when the hand differed. We focus on the relationship between stimulus and response independently from the side, meaning that ipsilateral conditions were concatenated in the same class across trials regardless of which hand was involved in the task; either left or right. The same logic was used for all the other conditions.

We compute classification between S-R conditions and Sor R-O conditions, depending on the alignment of the S-R data to the stimulus onset or response onset respectively. Both time-resolved and searchlight classification approaches were implemented. The first modality used all brain parcels as features for each time point; the second considered each brain parcel along with its 5 nearest spatial neighbours as features, estimating classification accuracy for each time point of a given time window, and then averaging across the time dimension.

### Multivariate regression

We implemented a multivariate linear regression analysis to predict reaction time based on brain activity patterns across both spatial (searchlight) and temporal dimensions, as for classification. Model performance was evaluated using R^2^ as the primary metric. A 10-fold cross-validation approach was employed to ensure robustness and generalizability, repeated twice to account for variability in the cross-validation splits.

Only S-R data aligned to the stimulus onset were used in this analysis, since the goal was to predict the response time relative to the stimuli. As for classification, conditions were aggregated based on the relationship between stimuli and response, hence regression was performed on ipsilateral and contralateral conditions independently.

To evaluate the statistical significance of the observed R^2^ values, we generated a null distribution through permutation testing with 2000 permutations. This approach involved randomly shuffling the reaction time labels (*y*) across trials while preserving the structure of the predictor data (*x*). The shuffling disrupts the true relationship between *x* and *y*, simulating a scenario where no meaningful association exists. For each permutation, the regression analysis was repeated, and the resulting R^2^ values were stored, creating a null distribution of R^2^ under the null hypothesis. The observed R^2^ values were then corrected for chance by subtracting the mean R^2^ of the null distribution. This correction accounts for spurious associations that might arise due to random fluctuations or noise in the data. The resulting corrected R^2^ values provide an estimate of the model’s performance beyond chance.

### Rolling expectation model

To investigate how trial-by-trial variability in stimulus onset timing reflects S-O and S-R association performance and was neurally encoded, we focused on fluctuations in ISIs. Although ISIs were randomly jittered between 2.5 and 3 seconds, we hypothesized that participants developed coarse temporal expectations based on the recent history of ISIs. To do so, we implied a simple rolling average model which estimates participants’ evolving expectations 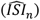 over time. For each trial *n*, the expected ISI was calculated as the mean of the preceding *ω* ISIs:

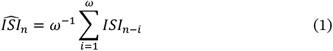

The parameter *ω*, referred to as the temporal window, determines the number of past trials used to generate each trial’s expectation. For early trials, where fewer than *ω* past trials were available, the expectation was computed using all available data (e.g., trial 3 used the mean of trials 1 and 2). The expectation value was further used to compute the prediction error *PE*_*n*_, defined as the difference between the actual ISI on trial *n* and the expected ISI:

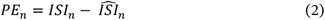

This model assumes that participants form temporal expectations based on the recent history of ISIs, reflecting a local, adaptive memory process. We also consider two additional configurations as baseline models: one assuming that participants hold fixed expectation based on the overall mean ISI across the entire session (*μ* model), while the other fixed the expected ISI at 0, thus equating the prediction error to the ISI on each trial *n* (ISI model).

Thus, for each participant we correlate each error trajectory with RTs using spearman correlation, pooling data across ipsilateral and contralateral conditions. Based on the hypothesis that a negative expectation error (i.e. ISI shorter than expected) would correspond to an extension of RTs compare to a positive error (i.e. ISI longer than expected) we selected the prediction error trajectory with the lowest median correlation value across participants.

The same model implied to generate this prediction error trajectory was also used to generate expectation error vector for S-O condition, which further served as comparative baseline allowing us to isolate neural signals specifically associated with the response-related encoding of temporal expectations.

### Neural encoding of prediction errors

Thus, we applied a multivariate linear regression analysis (same hyperparameter as in previous paragraph) to identify whether and when brain activity encodes this prediction error. For each participant and S-O or S-R conditions, time-locked neural data aligned to stimulus onset served as regressors and the trial-by-trial prediction error trajectory was used as the continuous outcome variable. For the S-R Ipsi condition we concatenated SLxRLx and SRxRRx trials, while for S-R Contra we concatenated SLxRRx and SLxRRx. To test whether expectation-related signals associated with the motor action were encoded strongly than passive stimulation, we compared S–O and S– R condition, enabling both temporal and spatial characterization of action-oriented neural encoding of temporal prediction errors.

Additionally, we performed a control analysis to ensure our results were not driven by the motor response. Thus, we averaged S-R trials time-locked to the motor response to generate a motor-evoked template. Then, we subtracted this template from all S-R trial aligned to each response time, obtaining a set of response-subtracted S-R trials. Thus, we repeated the same time-resolved multivariate regression analysis as above on these response-subtracted data. Furthermore, we tested the pattern generalization effects of these neural encoding mechanisms by performing cross-decoding analyses^24^ across sensory inputs and motor outputs. Specifically, for the sensory aligned condition we trained a model on SLxRLx and tested on SLxRRx to obtain a value of model performance (corrected R^2^) per time-point. Then, we inverted train and test data and repeated the cross-decoding procedure and average the two results, i.e. SLxRLx *→* SLxRRx and SLxRRx *→* SLxRLx. We then repeated the same procedure for SRxRLx and SRxRRx conditions and average the results across left and right sensory input side. In this way, the sensory input was held constant while the motor output was variable. Accordingly, for the motor aligned condition we trained a model on SLxRLx, tested it on SRxRLx, repeat the procedure inverting train and test sets and average the results. We then repeated the same procedure for SRxRRx and SLxRRx conditions and average the results across left and right motor output side. In this way, the motor output was held constant while the sensory input was variable.

### Statistical analysis

Behavioral data were tested for normality using Kolmogorov-Smirnov test. When normality assumption was violated, data were tested using a paired sample, two-tailed, Wilcoxon signed rank test. Neural data has been tested at the group level using cluster-based permutation statistics ^59^. First, one-tail paired-samples t-value were computed and selected when smaller than the critical value for α = 0.05. All selected samples were then clustered based on adjacency, either temporal (time series) or spatial (searchlight), and t-values within a cluster were summed to calculate cluster-level statistics. Finally, the p-value for all the selected clusters were calculated under the permutation distribution of the largest clusters, generated by means of the Monte Carlo method using all possible combinations (2^11^ = 2048 iterations). Spatial adjacency for statistics on searchlight analysis were derived from a topological neighbors structure, defined by the proximity of brain parcels. Effect sizes are reported using Cohen’s d.

## Supplementary material

**Supplementary Figure 1.**
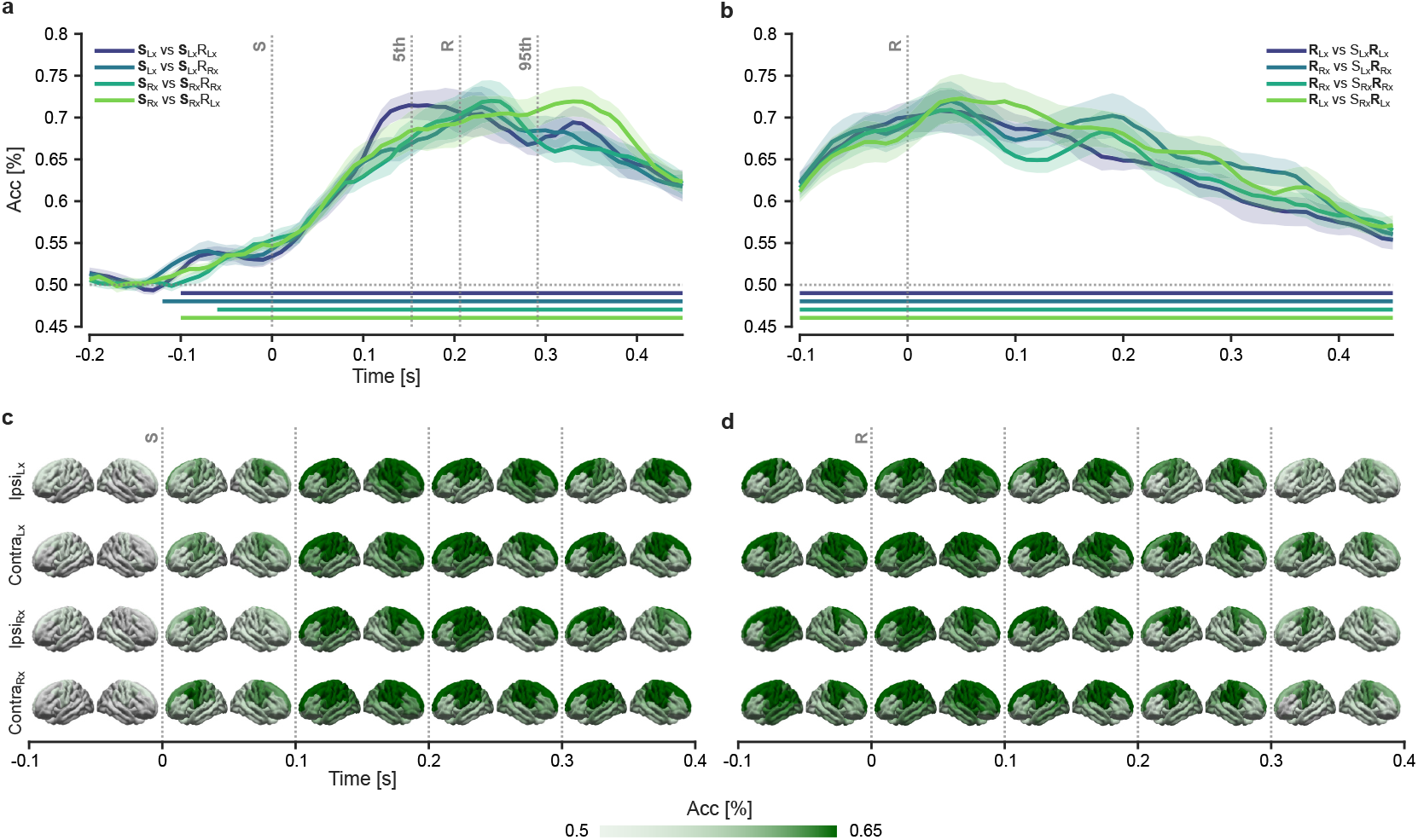
Decoding results for individual stimulus–response (S–R) conditions. Time-resolved multivariate decoding accuracy (Acc) discriminating S-R (all individual conditions) conditions from stimulus-only (S-O, panel **a**.) and response-only (R-O, panel **b**.) baseline conditions. Shaded areas indicate SEM, while horizontal bars denote statistical significance above chance level (cluster-based one-tailed permutation test, p < 0.05). Vertical dotted lines represent the stimulus onset (S), the response onset (R) and the 5^th^ and 95^th^ percentile of the temporal response onset distribution. Below, cortical surface maps from time-resolved searchlight decoding illustrate the spatial distribution of S–R decoding versus S-O (**c**.) and R-O (**d**.) across lateralized conditions. Statistical significance was determined by cluster based one-tail permutation test (p < 0.05).

**Supplementary Figure 2.**
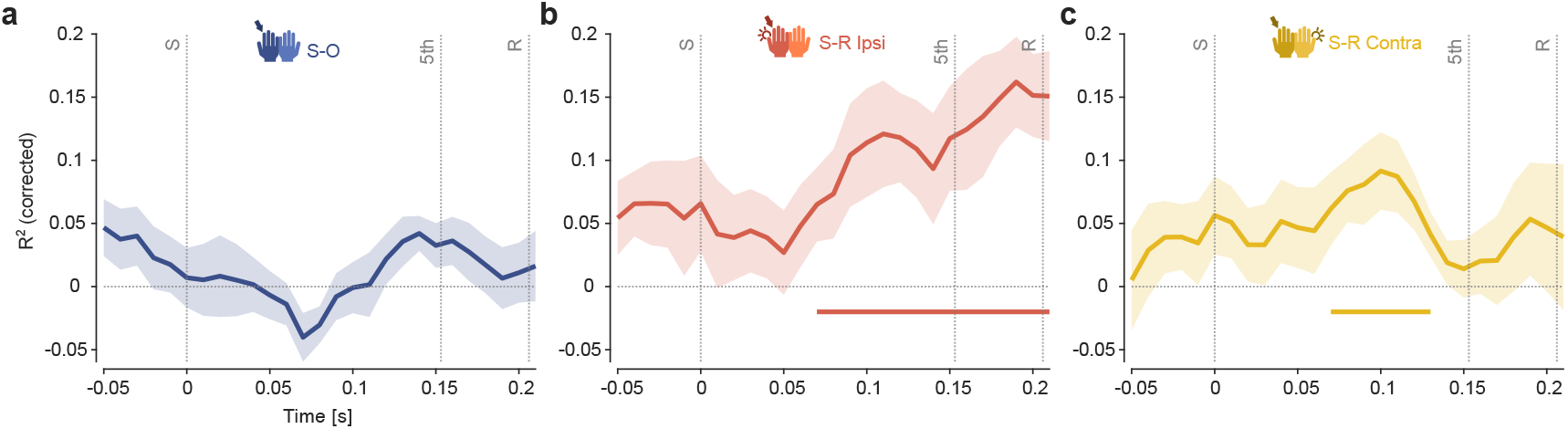
Neural encoding of temporal prediction errors on individual conditions. Lineplots showing the time-resolved multivariate regression (R^²^, corrected) model performance of peri-stimulus neural activity predicting trial-wise prediction error in S-O (**a**.), S-R Ipsi (**b**.) and S-R Contra (**c**.) conditions against chance level. Shaded areas represent SEM. Horizontal bars indicate statistical significance (cluster-based permutation test, p < 0.05). Vertical dotted lines represent the stimulus onset (S), the response onset (R) and the 5^th^ percentile of the temporal response onset distribution.

**Supplementary Figure 3.**
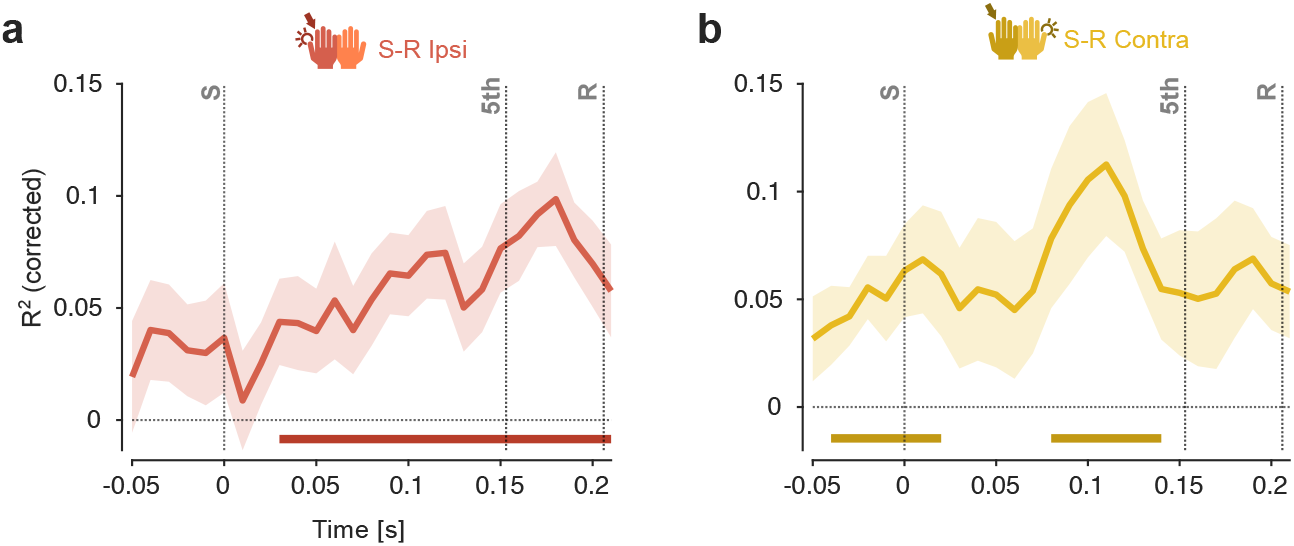
Neural encoding of temporal prediction errors on individual conditions on response-subtracted data. Lineplots showing the time-resolved multivariate regression (R^²^, corrected) model performance of peristimulus neural activity predicting trial-wise prediction error S-R Ipsi (**a**.) and S-R Contra (**b**.) conditions against chance level on response-subtracted data. Shaded areas represent SEM. Horizontal bars indicate statistical significance (cluster-based permutation test, p < 0.05). Vertical dotted lines represent the stimulus onset (S), the response onset (R) and the 5^th^ percentile of the temporal response onset distribution.

## References

1. Ackerley, R., Borich, M., Oddo, C. M. & Ionta, S. Insights and Perspectives on Sensory-Motor Integration and Rehabilitation. Multisensory Res. 29, 607–633 (2016).

2. Edwards, L. L., King, E. M., Buetefisch, C. M. & Borich, M. R. Putting the “Sensory” Into Sensorimotor Control: The Role of Sensorimotor Integration in Goal-Directed Hand Movements After Stroke. Front. Integr. Neurosci. 13, (2019).

3. Rao, R. P. & Ballard, D. H. Predictive coding in the visual cortex: a functional interpretation of some extra-classical receptive-field effects. Nat. Neurosci. 2, 79–87 (1999).

4. Friston, K. A theory of cortical responses. Philos. Trans. R. Soc. B Biol. Sci. 360, 815–836 (2005).

5. Kok, P., Mostert, P. & De Lange, F. P. Prior expectations induce prestimulus sensory templates. Proc. Natl. Acad. Sci. 114, 10473–10478 (2017).

6. Millidge, B., Seth, A. & Buckley, C. L. Predictive Coding: a Theoretical and Experimental Review. Preprint at http://arxiv.org/abs/2107.12979 (2022).

7. Greco, A. et al. Statistical Learning of Incidental Perceptual Regularities Induces Sensory Conditioned Cortical Responses. Biology 13, 576 (2024).

8. Greco, A., Moser, J., Preissl, H. & Siegel, M. Predictive learning shapes the representational geometry of the human brain. Nat. Commun. 15, 9670 (2024).

9. Bonetti, L. et al. Spatiotemporal brain hierarchies of auditory memory recognition and predictive coding. Nat. Commun. 15, 4313 (2024).

10. Greco, A., Rastelli, C., Bonetti, L., Braun, C. & Caria, A. Neural signatures of predictive coding underlying the acquisition of incidental sensory associations. 2025.05.16.654429 Preprint at 10.1101/2025.05.16.654429 (2025).

11. Greco, A., Rastelli, C., Ubaldi, A. & Riva, G. Immersive exposure to simulated visual hallucinations modulates high-level human cognition. Conscious. Cogn. 128, 103808 (2025).

12. Bonetti, L. et al. Shared and modality-specific brain networks underlying predictive coding of temporal sequences. 2025.07.16.665207 Preprint at 10.1101/2025.07.16.665207 (2025).

13. Badde, S., Myers, C. F., Yuval-Greenberg, S. & Carrasco, M. Oculomotor freezing reflects tactile temporal expectation and aids tactile perception. Nat. Commun. 11, 3341 (2020).

14. Ren, R. et al. Tactile temporal predictions: The influence of conditional probability. -Percept. 15, 20416695241264736 (2024).

15. Ren, R. et al. Electrophysiological evidence for the effect of tactile temporal prediction. Neuropsychologia 210, 109095 (2025).

16. Xiong, Z., Job, X. & Kilteni, K. Costs and benefits of temporal expectations on somatosensory perception and decision-making. Cognition 261, 106146 (2025).

17. Grabenhorst, M., Poeppel, D. & Michalareas, G. Neural signatures of temporal anticipation in human cortex represent event probability density. Nat. Commun. 16, 2602 (2025).

18. Den Ouden, H. E. M., Friston, K. J., Daw, N. D., McIntosh, A. R. & Stephan, K. E. A Dual Role for Prediction Error in Associative Learning. Cereb. Cortex 19, 1175–1185 (2009).

19. Stokes, M. G., Myers, N. E., Turnbull, J. & Nobre, A. C. Preferential encoding of behaviorally relevant predictions revealed by EEG. Front. Hum. Neurosci. 8, (2014).

20. Auksztulewicz, R., Friston, K. J. & Nobre, A. C. Task relevance modulates the behavioural and neural effects of sensory predictions. PLOS Biol. 15, e2003143 (2017).

21. Van Veen, B. D., Van Drongelen, W., Yuchtman, M. & Suzuki, A. Localization of brain electrical activity via linearly constrained minimum variance spatial filtering. IEEE Trans. Biomed. Eng. 44, 867–880 (1997).

22. Desikan, R. S. et al. An automated labeling system for subdividing the human cerebral cortex on MRI scans into gyral based regions of interest. NeuroImage 31, 968–980 (2006).

23. Cichy, R. M. & Pantazis, D. Multivariate pattern analysis of MEG and EEG: A comparison of representational structure in time and space. NeuroImage 158, 441–454 (2017).

24. Sandhaeger, F. & Siegel, M. Testing the generalization of neural representations. NeuroImage 278, 120258 (2023).

25. Gallivan, J. P., McLean, D. A., Flanagan, J. R. & Culham, J. C. Where One Hand Meets the Other: Limb-Specific and Action-Dependent Movement Plans Decoded from Preparatory Signals in Single Human Frontoparietal Brain Areas. J. Neurosci. 33, 1991–2008 (2013).

26. Ariani, G., Oosterhof, N. N. & Lingnau, A. Time-resolved decoding of planned delayed and immediate prehension movements. Cortex 99, 330–345 (2018).

27. Adam, J. J. et al. Rapid visuomotor preparation in the human brain: a functional MRI study. Cogn. Brain Res. 16, 1–10 (2003).

28. Ramayya, A. G., Buch, V., Richardson, A., Lucas, T. & Gold, J. I. Human response times are governed by dual anticipatory processes with distinct neural signatures. Commun. Biol. 8, 124 (2025).

29. Braun, C. et al. Functional Organization of Primary Somatosensory Cortex Depends on the Focus of Attention. NeuroImage 17, 1451–1458 (2002).

30. Dockstader, C., Cheyne, D. & Tannock, R. Cortical dynamics of selective attention to somatosensory events. NeuroImage 49, 1777–1785 (2010).

31. Miltner, W., Johnson, R., Braun, C. & Larbig, W. Somatosensory eventrelated potentials to painful and non-painful stimuli: effects of attention. Pain 38, 303–312 (1989).

32. Churchland, M. M., Yu, B. M., Ryu, S. I., Santhanam, G. & Shenoy, K. V. Neural Variability in Premotor Cortex Provides a Signature of Motor Preparation. J. Neurosci. 26, 3697–3712 (2006).

33. Duque, J., Labruna, L., Verset, S., Olivier, E. & Ivry, R. B. Dissociating the Role of Prefrontal and Premotor Cortices in Controlling Inhibitory Mechanisms during Motor Preparation. J. Neurosci. 32, 806–816 (2012).

34. Narayanan, N. S. & Laubach, M. Top-Down Control of Motor Cortex Ensembles by Dorsomedial Prefrontal Cortex. Neuron 52, 921–931 (2006).

35. Bolton, D. A. E. & Staines, W. R. Transient inhibition of the dorsolateral prefrontal cortex disrupts attention-based modulation of tactile stimuli at early stages of somatosensory processing. Neuropsychologia 49, 1928–1937 (2011).

36. Gogulski, J. et al. A Segregated Neural Pathway for Prefrontal TopDown Control of Tactile Discrimination. Cereb. Cortex 25, 161–166 (2015).

37. Staines, W. R., Graham, S. J., Black, S. E. & McIlroy, W. E. Task-Relevant Modulation of Contralateral and Ipsilateral Primary Somatosensory Cortex and the Role of a Prefrontal-Cortical Sensory Gating System. NeuroImage 15, 190–199 (2002).

38. Juravle, G. & Deubel, H. Action preparation enhances the processing of tactile targets. Exp. Brain Res. 198, 301–311 (2009).

39. St. John-Saaltink, E., Utzerath, C., Kok, P., Lau, H. C. & De Lange, F. P. Expectation suppression in early visual cortex depends on task set. PLoS One 10, e0131172 (2015).

40. Schlossmacher, I., Dellert, T., Pitts, M., Bruchmann, M. & Straube, T. Differential Effects of Awareness and Task Relevance on Early and Late ERPs in a No-Report Visual Oddball Paradigm. J. Neurosci. 40, 2906–2913 (2020).

41. Moskowitz, H. S. & Sussman, E. S. Sound category habituation requires task-relevant attention. Front. Neurosci. 17, (2023).

42. Duncan, D. H., van Moorselaar, D. & Theeuwes, J. Visual statistical learning requires attention. Psychon. Bull. Rev. (2024) doi:10.3758/s13423-024-02605-1.

43. Friston, K. et al. Active inference and learning. Neurosci. Biobehav. Rev. 68, 862–879 (2016).

44. Pezzulo, G., Rigoli, F. & Friston, K. J. Hierarchical active inference: a theory of motivated control. Trends Cogn. Sci. 22, 294–306 (2018).

45. Pezzulo, G., Parr, T. & Friston, K. The evolution of brain architectures for predictive coding and active inference. Philos. Trans. R. Soc. B Biol. Sci. 377, 20200531 (2022).

46. Ali, A., Ahmad, N., de Groot, E., van Gerven, M. A. J. & Kietzmann, T. C. Predictive coding is a consequence of energy efficiency in recurrent neural networks. Patterns 3, 100639 (2022).

47. Gibson, J. J. The Ecological Approach to Visual Perception. xiv, 332 (Houghton, Mifflin and Company, Boston, MA, US, 1979).

48. O’Regan, J. K. & Noë, A. A sensorimotor account of vision and visual consciousness. Behav. Brain Sci. 24, 939–973 (2001).

49. Varela, F. J., Rosch, E. & Thompson, E. The embodied mind. Embodied Mind Cogn. Sci. Hum. Exp. (1991).

50. Friston, K. The free-energy principle: a unified brain theory? Nat. Rev. Neurosci. 11, 127–138 (2010).

51. Kadiallah, A., Franklin, D. W. & Burdet, E. Generalization in Adaptation to Stable and Unstable Dynamics. PLOS ONE 7, e45075 (2012).

